# Predictive factors in identifying operative risks in cholecystectomies

**DOI:** 10.1101/694182

**Authors:** Murat Kanlioz, Ugur Ekici

**Author notes:** **Corresponding Author: Ugur Ekici, E mail**, **Adress**: Tahtakale mah. Ahmet Hamdi Tanpınar Sok. Avrupa Konutları 3 1-B Blok No:98 Avcılar /İstanbul, **Murat Kanlioz; Email**.

## Abstract

**Purpose:** This study aims to forecast findings showing the difficulty of operation in cholecystectomy through pre-operative examinations and reduce morbidity and mortality with the predictive data obtained.

**Materials and Methods:** In the preoperative period, the followings were measured in patients who will undergo cholecystectomy: C-reactive protein (CRP), erythrocyte sedimentation rate (ESR), WBC, Neutrophil ratio (NR), erythrocyte distribution range (RDW), aspartate aminotransferase (AST), alanine aminotransferase (ALT), alkaline phosphatase (ALP), gamma glutamyl transferase (GGT), total bilirubin (TB), direct bilirubin (DB). Following the preoperative ultrasound (USG), the patients were recorded in two groups as patients with “normal” and “increased” gallbladder wall thickness. Also, the patients were asked if they underwent ERCP and whether they received antibiotic treatment in the last 10 days due to their disease in the preoperative period. The appearance of the peroperative gallbladder was recorded in two groups as “has a normal appearance” or “edematous and/or adherent to peripheral tissues”. Whether or not there is a correlation between the preoperative findings and peroperative appearance was evaluated. The recordings and analyzes were made using SPSS statistics program. Correlation between the data were analyzed by Chi-square test. p<0.05 was considered significant.

**Results:** The study achieved statistically significant results for the correlation between the “gallbladder edema and/or adhesion to peripheral tissues” in the peroperative period and the following five parameters: increased WBC, increased NR, increased gallbladder wall thickness at USG, compulsory ERCP and receipt of antibiotic treatment for the disease in the last 10 days.(p<0,05).

**Conclusion:** Taking into consideration the presence, in the preoperative period, of some or all of the five criteria-namely, increased WBC, increased NR, increased gallbladder wall thickness at USG, receiving antibiotic treatment for the disease in the last 10 days and undergoing ERCP-in patients with cholelithiasis for whom cholecystectomy is envisaged would make it easier to estimate the degree of difficulty of the surgery and the possibility of encountering difficult and complicated cases.

## Introduction

In cholecystectomy, both preoperative and postoperative complications are still severe ^1, 2^. Predicting the factors that may lead up to complications and making a plan accordingly would both reduce the complications and strengthen the hand of surgeons in presenting preoperative risk analysis to patients. In this study, we intended to study whether there were preoperative indications for the picture that may be encountered during operation in patients scheduled for cholecystectomy. Despite not being the single factor determining morbidity and mortality, the condition of gallbladder as *normal* or *edematous and/or adherent to peripheral tissues* during surgery is one of the most important indicators. We also intended to analyze the correlation between the preoperative appearance of the gallbladder and the parameters we measured in the preoperative period. We thereby aimed to evaluate the predictability in the preoperative period of the possible picture that we may encounter during the operation. Hence, we tried to identify the predictive parameters by marking down the status of the preoperative gallbladder and its correlation with the preoperative clinical examination, laboratory findings and imaging results. In conclusion, we used to investigate whether a preoperative risk analysis is practicable thanks to the resulting possible risk parameters.

## Materials and Methods

The study included the first 200 patients who were admitted to our clinic due to gallbladder stone and underwent cholecystectomy. Along with the demographic data of the patients, the values CRP (0-0,5 mm/dL), ESR (0-20 mm/h), WBC (4,6-10,2 *10^3^/uL), NR (50-70%), RDW (11.6-17.2%), AST (5-30 U/L), ALT (0-55 U/L), ALP (40-150 U/L), GGT (12-64 U/L), TB (0.2-1.2 mg/dL), DB (0-0.5 mg/dL) measured in blood in the preoperative period were recorded, and the results were classified as *low, normal* and *high* as per their reference values. According to the preoperative ultrasonography findings, two groups were formed as with *increased* and *normal* gallbladder wall thickness. Whether the patients underwent a preoperative ERCP and whether they received antibiotic treatment in the last 10 days due to their disease are recorded. The preoperative findings were recorded as *has a normal appearance* or *edematous and/or adherent to peripheral tissues* according to the appearance of the gallbladder. The correlations between preoperative findings and preoperative findings were analyzed. The recordings and analyzes were made using the software SPSS. The groups were compared by Chi-square test. *p<0.05* was considered significant.

Our study complies with the principles of the Declaration of Helsinki. Local ethics committee approval was obtained.

## Results

The study included 200 patients, of whom 172 (86%) were female and 28 (14%) were male. The average age was 47.2±14,15 years, and the median age was 47 years.

Of 96 patients whose preoperative gallbladder *had a normal appearance*, 8 (8.33%) had increased WBC. Of 104 patients with a peroperative gallbladder *edematous and/or adherent to peripheral tissues*, 25 (24.03%) had increased WBC. The difference in-between was statistically significant (*p<0.001*). (Table 1)

**Table 1.** Correlation between peroperative appearance of the gallbladder and the preoperative parameters

| (n: number of patients) |  | Peroperative Appearance of Gallbladder |  |
| --- | --- | --- | --- |
|  |  | Has a normal appearance (n) | Edematous and/or adherent to peripheral tissues (n) |
| **Preoperative WBC** | Low |  |  |
|  | Normal | 6 | 4 |
|  | High | 82 | 75 |
|  | Total | 8 | 25 |
|  |  | 96 | 104 |
| **Preoperative NR** | Low |  |  |
|  | Normal | 6 | 15 |
|  | High | 70 | 66 |
|  | Total | 10 | 19 |
|  |  | 86 | 100 |
| **Preoperative ERCP** | Performed | 1 | 4 |
|  | Non-performed | 95 | 100 |
|  | Total | 96 | 104 |
| **Preoperative Antibiotic Use** | Yes |  |  |
|  | No | 21 | 38 |
|  | Total | 75 | 66 |
|  |  | 96 | 104 |
| **Wall Thickening in Preoperative USG** | Yes | 7 | 22 |
|  | No | 89 | 82 |
|  | Total | 96 | 104 |

Of 96 patients whose preoperative gallbladder *had a normal appearance*, 10 (11.62%) had increased NR. Of 104 patients with a preoperative gallbladder *edematous and/or adherent to peripheral tissues*, 19 (19%) had increased NR. The difference in-between was statistically significant (*p<0.01*). (Table 1). (Note: Since some of the patients in the NR group had no NR results, the number of patients in this group was 186.)

Of 96 patients whose peroperative gallbladder *had a normal appearance*, one (1.04%) stated to have undergone ERCP. Of 104 patients with a preoperative gallbladder *edematous and/or adherent to peripheral tissues*, 4 (3.84%) had not undergone ERCP. The difference in-between was statistically significant (*p<0.001*). (Table 1)

Of 96 patients whose peroperative gallbladder *had a normal appearance*, 21 (21.87%) stated to have received antibiotic treatment in the last 10 days prior to admission to the hospital. Of 104 patients with a peroperative gallbladder *edematous and/or adherent to peripheral tissues*, 38 (36.53%) stated to have received antibiotic treatment in the last 10 days before they were referred to the hospital. The difference in-between was statistically significant (*p<0.01*). (Table 1)

Of 96 patients whose preoperative gallbladder *had a normal appearance*, it was possible to determine the gallbladder wall thickness in 7 (7.29%) in the preoperative USG. Of 104 patients with a preoperative gallbladder *edematous and/or adherent to peripheral tissues*, the gallbladder wall thickening was determined in 22 (21.15%) in the preoperative USG. The difference in-between was statistically significant (*p<0.001*). (Table 1)

In the statistical analyses performed, no significant correlation was found between the preoperative appearance of gallbladder and the AST, ALT, ALP, GGT, TB, DB, CRP, SR and RDW measured in the preoperative period.

## Discussion

The mean age of patients in our study is 47.2±14.15 years. In their study, Pal et al. stated the mean age of patients as 58.5 years, where they studied the patients with high-risk acute cholecystitis ^3^. However, the mean age was reported to be 43,14±14,16 years in the study of Ahmed et al ^4^. The rate of female patients in our study was 86%. In the study of Brunee et al., the rate of female patients was reported to be 55.83% ^5^. We found a significant correlation between the preoperative gallbladder’s state of being *edematous and/or adherent to peripheral tissues* and the preoperative WBC rate. In their study, Brunee et al. reported that high WBC values increased operative complication and hospitalization period ^5^. Portinari et al. reported in their study that increased WBC, increased CRP and gallbladder wall thickening at USG were among the most important criteria in the need for emergency surgery in cholelithiasis ^6^. We found no statistical significance between increased CRP and the peroperative gallbladder’s state of being *edematous and/or adherent to peripheral tissue*. However, in their studies, *Portinari et al., Lee et al.* and *Oneo et al.* reported to have found a significant correlation between CRP and the patients with complicated cholelithiasis ^6-8^. We found a statistical significance between the peroperative gallbladder’s state of being *edematous and/or adherent to peripheral tissues* and the preoperative NR rate. In their study, Bourgouin et al. also reported to have found a significant correlation between NR and complicated cholecystectomies ^9^. We also found a statistical significance between those who underwent ERCP and the peroperative gallbladder’s state of being *edematous and/or adherent to peripheral tissues*. In all their studies, *Donkervoort et al., Lee et al.* and *Spens et al.* reported that having undergone ERCP is an important finding that the operative risk might be increased ^1, 7, 10^. Our study also revealed a statistical significance between those who received antibiotic treatment within the last 10 days prior to admission to the hospital and the peroperative gallbladder’s state of being *edematous and/or adherent to peripheral tissues*. This parameter was realized in a quality we predicted before conducting the study.

We found a significant correlation between the peroperative gallbladder’s state of being *edematous and/or adherent to peripheral tissues* and the gallbladder wall thickening in preoperative USG. In their study, Izquierdo et al. reported to have found that having a gallbladder wall thickness of 6 mm and above at USG had a 87.5% sensitivity in predicting return to open surgery ^11^.

The fact that, in the statistical analyses performed, no significant correlation was found between the preoperative appearance of gallbladder and the AST, ALT, ALP, GGT, TB, DB, CRP, ESR and RDW measured in the preoperative period contradicted our pre-study predictions.

## Conclusion

Taking into consideration, prior to operation, the presence, in the preoperative period, of some or all of the five criteria-namely, increased WBC, increased NR, increased gallbladder wall thickness at USG, receiving antibiotic treatment for the disease in the last 10 days and undergoing ERCP-is envisaged would make it easier to estimate the degree of difficulty of the surgery and the possibility of encountering difficult and complicated cases. Scoring the five parameters on which we found a significant correlation by giving one point for each, we may say that the operative risk may increase in proportion to the increase in score.

